# Development and Evaluation of a Machine Learning Recursive Partitioning Decision Tree Algorithm to Optimize Urinalysis Parameters to Predict Urine Culture Positivity

**DOI:** 10.1101/2023.03.10.532117

**Authors:** Jansen N. Seheult, Michelle N. Stram, Lydia Contis, Raymond E. Pontzer, Stephanie Hardy, William Wertz, Carla M. Baxter, Michael Ondras, Paula L. Kip, Graham M. Snyder, A. William Pasculle

## Abstract

PittUDT, a recursive partitioning decision tree algorithm for predicting urine culture (UC) positivity based on macroscopic and microscopic urinalysis (UA) parameters, may significantly decrease the number of unnecessary UC. The reflex algorithm derivation included 38,361 paired UA and UC cases; the average patient age was 57.4 years and 70% of samples were from female patients. Receiver operating characteristic (ROC) analysis identified urine WBCs, leukocyte esterase, and bacteria as the best predictors of UC positivity, with area under the ROC curve of 0.79, 0.78, and 0.77, respectively. The test data comprised 9,773 cases, of which 2,571 (26.3%) had a positive UC result. Using the held-out test data set, the PittUDT algorithm met the pre-specified targets of a 30-60% total negative proportion (true negative plus false negative predictions) with a negative predictive value above 90%. These data show that a rule-based algorithm trained on UA data has adequate predictive ability for triaging urine specimens by identifying low risk urine specimens, which are unlikely to grow pathogenic organisms, with a false-negative proportion under 5%. Our study demonstrates the feasibility and performance of using a supervised machine learning approach to develop a human readable machine learning model to optimize UA parameters in a reflex protocol that can be deployed across multiple hospital sites and settings. Our work supports a UA reflex to UC protocol that has the capacity to optimize UA parameters to predict UC positivity and improve antimicrobial stewardship and UC utilization, a potential avenue for cost savings.

## INTRODUCTION

The diagnosis of urinary tract infections (UTIs) from history alone is challenging due to the nonspecific signs and symptoms that can accompany these infections. Various combinations of macroscopic and microscopic urinalysis (UA) parameters in the setting of a clinical history and physical examination are used for making a presumptive diagnosis of UTI.^1^ When used in patient populations with undifferentiated abdominal pain or even among uncomplicated patients with typical UTI symptoms, the yield of routine urine cultures (UC) is quite low.^2–4^ Protocols which encourage more selective use of UC may improve antimicrobial stewardship and UC utilization, a potential avenue for cost savings. Reflex testing refers to an algorithm, which uses preliminary screening tests to inform decisions about the need for further tests. It has been applied in many settings in laboratory medicine, including protein electrophoresis and immunofixation, celiac disease serology, and thyroid function tests. Protocols for the implementation of UA with automated reflex to UC have been investigated in several settings, including in urology clinics, family practice outpatient clinics, and in the emergency department (ED).^5,6^ The most commonly used UA parameters are the presence of bacteria, white blood cells (WBCs), nitrites, or leukocyte esterase. Jones et al. have shown that the use of such a reflex protocol can reduce the number of UC performed in an ED setting by up to 39% while only missing 11 of 314 positive UC (3.5%).^7^

Traditional algorithms for UC reflex protocols have relied on univariate receiver operating characteristic curve analysis or multivariate (usually logistic) regression analysis using variables which had been categorized based on expert opinion and clinical judgment. The conventional approach of first performing univariate analyses and including in the multivariable model only those variables which meet a predetermined (usually liberal) significance level in univariate analyses ignores the fact that the significance of a predictor is affected by other variables. Furthermore, supervised machine learning algorithms offer an alternative approach to the automatic identification of useful rules with minimum human input. Classification and regression tree algorithms produce an output that is human-readable and simple to implement within the current laboratory information system infrastructure. These algorithms also permit differential weighting of false positives and false negatives in order to address class imbalances, i.e., where one class or category is under-represented in the data set. Differential weighting also allows the development of an algorithm that stresses negative predictive value. These decision tree algorithms have the potential to produce prediction tools that are more accurate compared to traditional methods.

In this study, we trained and validated a recursive partitioning decision tree algorithm for predicting UC positivity based on macroscopic and microscopic UA features. We also describe implementation of this UA reflex protocol under which UC orders are canceled if an accompanying automated UA does not meet pre-specified criteria and the method for performance monitoring after protocol implementation.

## MATERIALS AND METHODS

### Setting and study population

A data set of UA and UC results performed between January 1, 2017 and December 31, 2017 from adult patients (≥ 18 years of age) in non-maternity inpatient and outpatient units at five hospitals of the UPMC (University of Pittsburgh Medical Center) academic healthcare system were extracted from the electronic health record and laboratory information system (LIS). Only patients with a UA and UC performed within 24 hours of each other were included in this analysis.

### Characterization of laboratory procedures and electronic reporting

The UA and UC standard operating procedures (SOPs) were reviewed for the 5 hospital sites to assess for differences in UA and UC parameter reporting, as well as differences in the how the Urinalysis WAM (Sysmex, Lincolnshire, IL) middleware rules transformed raw counts into binned ordinal values at each site. For the 5 hospital sites, the LIS Sunquest (Sunquest, Tucson, AZ) was queried to determine all order codes used for UA and UC, the names of all urinalysis-associated specimen types, and the LIS test result codes (value IDs) for each of the UA parameters.

### Microbiologic methods

All sites performed the microscopic UA using the Sysmex UF-1000i (Sysmex, Lincolnshire, IL), which performs identification and quantification via flow cytometry, and the Clinitek Atlas or Clinitek Novus (Clinitek, Ramsey, MN) for the macroscopic UA; and manual microscopy was performed when indicated according to each laboratory’s SOPs. UC were performed in each hospital’s microbiology laboratory using standard methodologies. The UA parameters that were studied included presence of WBCs, bacteria, red blood cells (RBCs), leukocyte esterase, nitrate, specific gravity, pH, character, protein, and blood. UC were considered positive if at least 10,000 colony-forming units (CFU)/mL of likely urinary pathogens were present, including Gram-negative rods, *Enterococcus* spp., *Staphylococcus saprophyticus*, *Staphylococcus aureus*, group B streptococci, and *Aerococcus*.^6^ UC that grew *Lactobacillus*, *Gardnerella vaginalis*, and diphtheroids alone or mixed flora were classified as negative.

### Reflex algorithm derivation

A three-step algorithm was created: Macroscopic UA to Microscopic UA to UC. The first decision tree (Macroscopic to Microscopic UA reflex) was trained using parameters obtained from the macroscopic UA: leukocyte esterase, nitrate, specific gravity, pH, character, protein, and blood. All observations that were classified as high-risk by the first decision tree were then used to train or validate the second decision tree (Microscopic UA to UC reflex), which included parameters obtained from the microscopic UA: WBCs, bacteria, and RBCs. Age, gender, hospital location, and specimen type were not used as predictors since they may not be consistently captured in LIS at all sites in the healthcare system.

### Receiver operating characteristic analysis (ROC)

ROC curves were plotted to compare the accuracy of macroscopic and microscopic UA parameters for predicting positive UC results.

### Recursive partitioning algorithm

The “rpart”^8^ recursive partitioning algorithm was implemented using “caret”^9^ in R version 3.4.2 (Comprehensive R Archive Network)^10^ and has been extensively described previously.^11^ Briefly, the decision tree was built by first finding the single variable which best separated the data into two subgroups known as daughter nodes; after the data were separated, this process was repeated on each subgroup until the subgroups reached a minimum size or until there could be no further improvement. At each node or decision point, sample partitioning was based on the predictor variable which maximized the goodness-of-split, i.e., created subgroups were more homogenous or ‘purer’ than the data in the original parent group. Variable importance ranking was based on the Gini Index, a measure of node impurity that favors larger partitions compared with other split criteria.^5^ Ten-fold cross-validation was used to give an internal estimate of misclassification by the decision tree and to avoid overfitting. The balanced accuracy was used as the training metric. The decision tree was pruned, i.e., potentially unnecessary nodes and branches were removed, by specifying the complexity parameter at which the balanced accuracy was highest.

Data from January 1, 2017 to September 30, 2017 were used to train and test the decision tree; the original data were randomly divided into training and test datasets in a 2:1 ratio. The decision trees were trained using the training dataset then the performance of the optimized decision trees was subsequently determined using the test dataset.

### Training of decision trees

Training parameters were optimized to achieve a total negative proportion (TNP: % true negatives + false negatives as a proportion of all cases) between 30% and 60% and a false negative proportion (FNP: % false negatives as a proportion of all cases) less than 5.0%. For the Macroscopic to Microscopic UA reflex, the training data set comprised 19,511 unique paired UA and UC results with complete data for all predictor variables. The minimum node size before a split could be attempted was set as 500 observations and the minimum bucket size for daughter nodes was set as 250 observations. A false negative was weighted eight times higher than a false positive in order to optimize the negative predictive value.

For the Microscopic UA to UC reflex, a separate decision tree was trained using the 11,136 observations from the original training data set above that were predicted as being high-risk or positive. The minimum node size before a split could be attempted was set as 500 observations and the minimum bucket size for daughter nodes was set as 250 observations. A false negative was weighted eight times higher than a false positive in order to optimize the negative predictive value.

Both decision trees were then combined into a stepwise UA decision tree algorithm called PittUDT (Macroscopic UA to Microscopic UA to UC).

### Performance evaluation using test data set

The test data set comprised 9,773 unique paired UA and UC results with complete data for all predictor variables. The PittUDT algorithm was applied in a stepwise fashion for classification of observations based on their Macroscopic UA and/or Microscopic UA results.

A true positive means that the algorithm correctly predicted that the UC was positive, and a true negative means that the algorithm correctly predicted that the UC was negative. A false positive prediction indicates that the model predicted that the UC was positive when in fact it was negative. A false negative prediction means that the model predicted that the UC was negative, when in fact it was positive.

Sensitivity (recall), specificity, positive predictive value (precision), and negative predictive value were calculated as previously described.^12^ Naïve accuracy was calculated as the proportion of all cases that were correctly classified. Balanced accuracy was calculated as the average of the proportions of each individual class that were correctly classified (i.e., the average of the sensitivity and specificity). The 95% confidence intervals (CI) for test performance metrics were calculated using the exact binomial method. The balanced accuracy was compared among different prediction tools non-parametrically with a chi-squared test.

The performance targets were a total negative proportion (TNP: % true negatives + false negatives as a proportion of all cases) between 30-60% with a negative predictive value above 90% and a false-negative proportion below 5% for each site.

### Deployment of the Reflex Protocol

Deployment of the Urine Infection Testing algorithm across the 5 sites, began in January 2021 ending in September 2022. The protocols for algorithm development and the Urine Infection Testing program were ethically reviewed and approved by the University of Pittsburgh’s Quality Improvement Committee (projects 1203 and 1722). Implementation of this algorithm included standardization of UA reporting at the 5 hospital sites, standardization of the middleware rules, generation of LIS rules using the decision tree algorithm, and development of a QC program for culturing a random selection of urine samples, predicted to be low risk by the algorithm, every month for performance qualification. Prior to rolling out the algorithm, a set of rules was developed for the physician order entry module of the hospital information system. Persons ordering UA with reflex to culture were required to designate by means of check boxes that the patient was not pregnant, not immunosuppressed, not undergoing urologic surgery, or that the specimen did not come from outside the bladder (e.g., nephrostomy). These samples were cultured regardless of the UA results.

### Quality control simulation

Data from all samples tested between October 1, 2017 and December 31, 2017 were used to simulate implementation of the validated algorithm at all five sites. A QC program was also simulated based on a two-stage hypergeometric sampling plan; unlike the binomial approach, the hypergeometric approach assumes that a finite number of urine samples will be submitted for testing during a monthly QC period.^13,14^ The estimated number of urine samples tested during a given 4-week QC period per site was calculated based on the average test volume per month at each site in the training data. The tolerance limit for the QC false negative proportion was set at 20%.

In the first stage of the sampling plan, up to four QC false negative results *(m* = 4) were permitted per QC period at each site. For each fixed population size, *N*, and number of allowed failures, *m*, the first stage sample size was calculated as the minimum number *n_1_* such the probability of observing *m* or fewer process failures in a sample of size *n_1_* from a population of size *N* is at most 10% when the true QC false negative proportion (or process failure rate) is 20% or more. If no more than one QC false negative result was observed among these *n_1_* QC samples in a 4-week period at a given site, the process was considered to be in control and no further QC testing was required during that period. For logistic reasons, the *n_1_* QC samples were divided evenly among the four weeks in each QC period and QC samples were chosen at random from the samples submitted for testing in that week. If there were more than 4 QC false negative results in the initial QC sample, a second QC sample was submitted for testing in that QC period.

The second stage sample size was calculated as the minimum number *n_2_* such that the probability of observing (i) one failure among the first *n_1_* samples at stage one and (ii) no more than one failure among the *n_2_* samples at stage two was less than 0.10 minus the probability of observing *m* or fewer process failures in the first stage when the true QC false negative proportion was at least 20%.

Conformance was demonstrated if no more than one additional process failure was observed in the second sample. If more than one additional process failure is observed, then QC testing has failed for that month and requires further investigation into algorithm performance and consideration of expanding the QC sample size to encompass all collected specimens in the upcoming month.

In addition to the QC program simulation, the total negative proportion and overall false negative proportion for all samples at each site were also determined. The tolerance limits for the total negative rate were 20 to 60%. The QC program was simulated 1,000 times, each time drawing a random sample of observations or cases from the QC data period; the proportion of simulations for each site with QC testing failures was determined to evaluate the robustness of the QC program.

### Quality control during real-world use

QC testing to ensure proper functioning of the algorithm at each site was conducted incrementally as the Urine Infection Testing implementation was rolled out. A total of three randomly selected patient specimens predicted by the PittUDT Urine Infection Testing algorithm as low risk were cultured per day. None of the specimens which underwent QC testing originally qualified for culture, either due to negative macroscopic or microscopic analysis described above in microbiologic methods. Testing was performed by trained microbiology staff members in a biological safety cabinet. The following guidelines were used to conduct quality control:

1. Urine specimens were planted to Blood, MacConkey, and CNA agars, and incubated in ambient air at 35°C.
2. Cultures were reviewed for clinically significant growth at 24 and 48 hours, post-incubation.

a. Clinically significant growth includes >10,000 cfu/mL of the following organisms:

i. Any Gram-negative rod
ii. β-hemolytic *Streptococcus* species
iii. *Enterococcus* species
iv. *Staphylococcus aureus*
v. Yeast or other fungi
b. Cultures demonstrating clinically significant growth were reviewed by the Infectious Disease team, to determine if it represents a true failure of the Urine Infection Testing process.
3. Culturing continued for 20 consecutive days.

a. Growth of one or less clinically significant organisms during the 20-day period demonstrated correct functioning of the Urine Infection Testing program algorithm.

## RESULTS

### Data cleansing prior to algorithm development

To generate algorithm rules using the historic data from the 5 sites, the data underwent cleansing to create a single unified dataset with values that were comparable across sites. Evaluation of the SOPs, middleware rules, urinalysis-associated specimen types, UA and UC related test codes, and the LIS test result codes (value IDs) for each of the UA/UC parameters revealed differences across sites which needed to be comprehensively evaluated, documented, and accounted for before algorithm development. To classify the location type of the patient when the UA/UC was performed, approximately 380 location codes for the 5 hospitals were identified, reviewed, and divided into location categories (**Table 1**); maternity-specific locations were excluded. All LIS test result codes were identified to retrieve results for all analytes in the UA and UC (for most analytes that make up the UA, each site had its own individual analyte code). Approximately 14 UA and 5 UC test order codes were identified among the 5 sites. Thirty-five values for specimen types were identified and manually reviewed and condensed into the categories represented in the data (**Table 2**) following data cleansing (e.g., identification of duplicate specimen types with non-meaningful differences due to the lack of uniform naming practices of each site such as “urine” and “urn” and free text values that did not contribute meaningful additional information such as “scath” free texted as “urine scath”).

**TABLE 1.** Patient demographics, location of specimen collection, and specimen source for all cases by hospital

| Characteristic | Hospital 1 | Hospital 2 | Hospital 3 | Hospital 4 | Hospital 5 | Overall |
| --- | --- | --- | --- | --- | --- | --- |
| Hospital bed count | 495 | 380 | 437 | 795 | 520 | - |
| Number of cases | 3,356 | 8,342 | 11,093 | 9,448 | 6,122 | 38,361 |
| Age, mean ± SD (range) | 60.1 ± 20.1<br>(18 - 116) | 41.7 ± 19.7<br>(18 - 99) | 63.2 ± 21.5<br>(18 - 108) | 58.6 ± 18.2<br>(18 - 100) | 65.0 ± 17.4<br>(18 - 104) | 57.4 ± 21.4<br>(18 - 116) |
| Gender, Male %: Female % | 1,267: 2,089<br>(37.8: 62.3) | 468: 7,874<br>(5.6: 94.4) | 3,175: 7,918<br>(28.6: 71.4) | 4,142: 5,306<br>(43.8: 56.2) | 2,487: 3,635<br>(40.6: 59.4) | 11,539: 26,822<br>(30.1: 69.9) |
| Location of specimen collection, n (%) |  |  |  |  |  |  |
| Emergency department | 1,476 (44.0) | 2,370 (28.4) | 7,175 (64.7) | 2,127 (22.5) | 1,871 (30.6) | 15,019 (39.2) |
| Intensive care unit | 120 (1.4) | 248 (2.2) | 699 (7.4) | 627 (10.2) | 1,890 (4.9) | 1,880 (4.9) |
| Inpatient floor unit | 851 (10.2) | 1,442 (13) | 4,016 (42.5) | 2,495 (40.8) | 9,679 (25.2) | 9,634 (25.2) |
| Outpatient | 5,001 (60) | 2,228 (20.1) | 2,431 (25.7) | 1,129 (18.4) | 11,598 (30.2) | 11,549 (30.3) |
| Other | 0 (0) | 0 (0) | 175 (1.9) | 0 (0) | 175 (0.5) | 175 (0.5) |
| Specimen source, n (%) |  |  |  |  |  |  |
| Clean catch | 3,081 (91.8) | 7,968 (95.5) | 10,781 (97.2) | 8,347 (88.4) | 5,304 (86.6) | 35,481 (92.5) |
| Foley | 92 (1.1) | 288 (2.6) | 944 (10) | 781 (12.8) | 2,337 (6.1) | 2,330 (6.1) |
| Loop nephrostomy | 2 (0) | 0 (0) | 5 (0.1) | 4 (0.1) | 18 (0.1) | 18 (0.1) |
| Nephrostomy | 23 (0.3) | 4 (0) | 38 (0.4) | 20 (0.3) | 100 (0.3) | 100 (0.3) |
| Straight catheter | 248 (3) | 13 (0.1) | 80 (0.9) | 3 (0.1) | 344 (0.9) | 344 (0.9) |
| Suprapubic catheter | 9 (0.1) | 7 (0.1) | 34 (0.4) | 10 (0.2) | 81 (0.2) | 81 (0.2) |
| Time difference between UA and UC, hours | 0 | 0 | 0 | 0 | 0 | 0 |
| median (5<sup>th</sup> to 95<sup>th</sup> percentiles) | (0 to 8.0; | (-0.7 to 2.0; | (0 to 0.5; | (-1.1 to 11.4; | (-0.1 to 8.3; | (-0.2 to 5.3; |
| (range) | -24.0 to 24.0) | -23.6 to 24.0) | -23.2 to 24.0) | -24.0 to 24.0) | -23.8 to 24.0) | -24.0 to 24.0) |

SD, standard deviation; UA, urinalysis; UC, urine culture.

**TABLE 2.** Summary of test data set decision tree performance by patient demographics and specimen source

| Demographics and Specimen Source |  |  |  |  |  |  |  |  |  |  |  |
| --- | --- | --- | --- | --- | --- | --- | --- | --- | --- | --- | --- |
| Performance Metric<br>(n = 9,773) | Overall | Male | Female | Age < 60<br>years | Age ≥ 60<br>years | Clean<br>Catch | Foley<br>Catheter | Loop<br>Neph | Neph | Straight<br>Catheter | Suprapubic<br>Catheter |
| Requires<br>micro UA,<br>n (%) | 5,673 (58) | 1,383 (46.7) | 4,290 (63) | 2,486 (52) | 3,187 (63.9) | 5,191 (57.4) | 389 (65.7) | 5 (100) | 19 (82.6) | 52 (55.9) | 17 (100) |
| Requires UC,<br>n (%) | 4,579 (46.9) | 1,101 (37.2) | 3,478 (51.1) | 2,023 (42.3) | 2,556 (51.2) | 4,158 (46) | 336 (56.8) | 5 (100) | 18 (78.3) | 46 (49.5) | 16 (94.1) |
| TP,<br>n (%) | 2,119 (21.7) | 485 (16.4) | 1,634 (24) | 758 (15.8) | 1,361 (27.3) | 1,922 (21.3) | 139 (23.5) | 5 (100) | 13 (56.5) | 29 (31.2) | 11 (64.7) |
| TN,<br>n (%) | 4,742 (48.5) | 1,772 (59.8) | 2,970 (43.6) | 2,535 (53) | 2,207 (44.2) | 4,459 (49.3) | 235 (39.7) | 4 (0) | 43 (17.4) | 1 (46.2) | (5.9) |
| FP,<br>n (%) | 2,460 (25.2) | 616 (20.8) | 1,844 (27.1) | 1,265 (26.4) | 1,195 (24) | 2,236 (24.7) | 197 (33.3) | 5 (0) | 17 (21.7) | 5 (18.3) | (29.4) |
| FN,<br>n (%) | 452 (4.6) | 88 (3) | 364 (5.3) | 226 (4.7) | 226 (4.5) | 426 (4.7) | 21 (3.5) | 0 (0) | 1 (4.3) | 4 (4.3) | 0 (0) |
| TNP, % | 53.1 | 62.8 | 48.9 | 57.7 | 48.8 | 54.0 | 43.2 | 0.0 | 21.7 | 50.5 | 5.9 |
| Sensitivity,<br>% (95% CI) | 82.4 (80.9 - 83.9) | 84.6 (81.4 - 87.5) | 81.8 (80 - 83.5) | 77 (74.3 - 79.6) | 85.8 (83.9 - 87.4) | 81.9 (80.2 - 83.4) | 86.9 (80.6 - 91.7) | 100 (47.8 - 100) | 92.9 (66.1 - 99.8) | 87.9 (71.8 - 96.6) | 100 (71.5 - 100) |
| Specificity,<br>% (95% CI) | 65.8 (64.7 - 66.9) | 74.2 (72.4 - 75.9) | 61.7 (60.3 - 63.1) | 66.7 (65.2 - 68.2) | 64.9 (63.2 - 66.5) | 66.6 (65.5 - 67.7) | 54.4 (49.6 - 59.2) | - | 44.4 (13.7 - 78.8) | 71.7 (58.6 - 82.5) | 16.7 (0.4 - 64.1) |
| PPV, %<br>(95% CI) | 46.3 (44.8 - 47.7) | 44.1 (41.1 - 47) | 47 (45.3 - 48.7) | 37.5 (35.4 - 39.6) | 53.2 (51.3 - 55.2) | 46.2 (44.7 - 47.8) | 41.4 (36.1 - 46.8) | 100 (47.8 - 100) | 72.2 (46.5 - 90.3) | 63 (47.5 - 76.8) | 68.8 (41.3 - 89) |
| NPV, %<br>(95% CI) | 91.3 (90.5 - 92.1) | 95.3 (94.2 - 96.2) | 89.1 (88 - 90.1) | 91.8 (90.7 - 92.8) | 90.7 (89.5 - 91.8) | 91.3 (90.5 - 92.1) | 91.8 (87.7 - 94.9) | - | 80 (28.4 - 99.5) | 91.5 (79.6 - 97.6) | 100 (2.5 - 100) |
| NA, %<br>(95% CI) | 70.2 (69.3 - 71.1) | 76.2 (74.6 - 77.7) | 67.6 (66.5 - 68.7) | 68.8 (67.5 - 70.1) | 71.5 (70.2 - 72.8) | 70.6 (69.6 - 71.5) | 63.2 (59.1 - 67.1) | 100 (47.8 - 100) | 73.9 (51.6 - 89.8) | 77.4 (67.6 - 85.4) | 70.6 (44 - 89.7) |
| BA, %<br>(95% CI) | 74.1 (70.3 - 72) | 79.4 (74.7 - 77.9) | 71.7 (67.7 - 69.8) | 71.9 (67.1 - 69.9) | 75.3 (71.8 - 74) | 74.2 (70.4 - 72.2) | 70.6 (63.7 - 70.1) | - | 68.7 (50.3 - 83) | 79.8 (69.2 - 85) | 58.3 (42 - 74.7) |

BA, balanced accuracy; CI, confidence interval; FN, false negative; FNP, false negative proportion; FP, false positive; Loop Neph, loop
nephrostomy; NA, naïve accuracy; Neph, nephrostomy; NPV, negative predictive value; PPV, positive predictive value; TN, true negative;
TNP, total predicted negative proportion; TP, true positive; micro UA, microscopic urinalysis; UC, urine culture.

Although all sites were using the Sysmex UF-1000i and the Clinitek Atlas or Novus, evaluation of the middleware rules for each site revealed differences in the way raw values were being binned into ordinal values. Using the middleware rules, the range of values for each bin of UA results was identified to establish comparable values between sites. Values for the UA parameters were also condensed into groups sharing an equivalent result where different reporting terms were used (e.g., leukocyte esterase “negative, trace, small, moderate, large” and “negative, trace, P1, P2, P3”). In all cases, the numerical values underlying the ordinal bin names were reviewed to ensure that the same values were being appropriately categorized together. This data cleansing and compiling process was repeated for each analyte for all UA and UC laboratory results.

### Reflex algorithm derivation

This derivation of the reflex algorithm included 38,361 paired UA and UC cases; the average patient age was 57.4 years and 70% of samples were from female patients (**Table 1**). The majority of specimens originated in the ED (39.2%), outpatient units (30.3%), and inpatient units (25.2%), and more than 98% of specimens were either clean catch (92.5%) or Foley catheter (6.1%) specimens. The sample collection time for the UA and UC specimens was within 5.3 hours for 95% of samples. Summary data for results of macroscopic and microscopic UA for all cases are provided in the supplemental materials (**Tables S1** and **S2**).

The training data set comprised 19,511 cases, of which 5,223 (26.8%) had a positive UC result. ROC analysis using the training data identified urine WBCs, leukocyte esterase, and bacteria as the best predictors of UC positivity, with area under the ROC curve (AUC) of 0.79, 0.78, and 0.77, respectively; the poorest predictors of UC positivity were specific gravity and pH, with AUC of 0.48 and 0.51, respectively (**Figure 1**). The optimized recursive partitioning decision tree is presented in **Figure 2**; the two-step algorithm included leukocyte esterase and nitrate for predicting whether microscopic UA should be performed and WBC and bacteria, in combination with the above two macroscopic parameters for predicting UC positivity.

**FIGURE 1:**
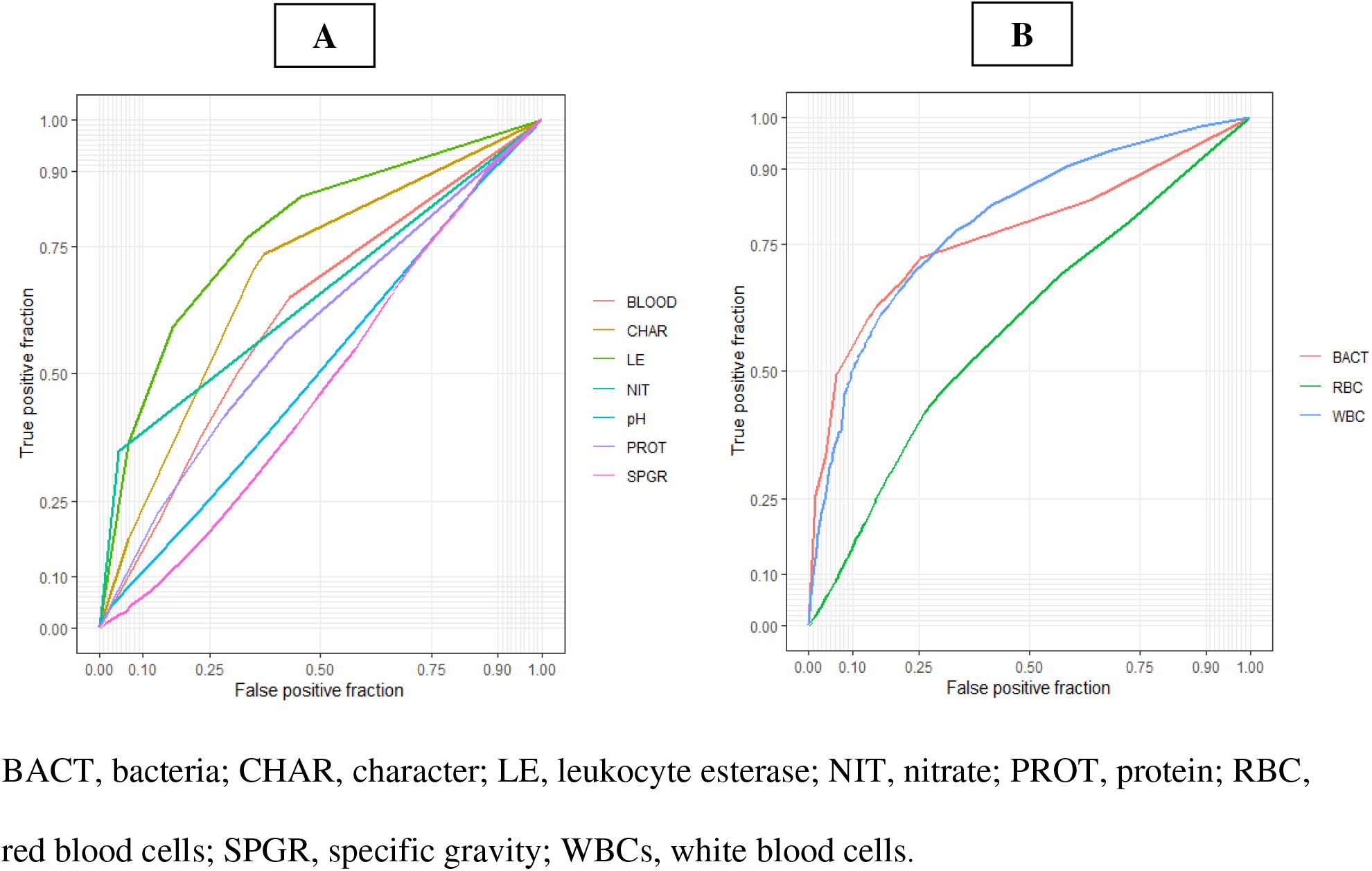
Receiver operating characteristic curves for prediction of urine culture positivity using (A) macroscopic urinalysis parameters and (B) microscopic urinalysis parameters. The area under the receiver operating characteristic curve was highest for white blood cells, leukocyte esterase, and bacteria.

**FIGURE 2:**
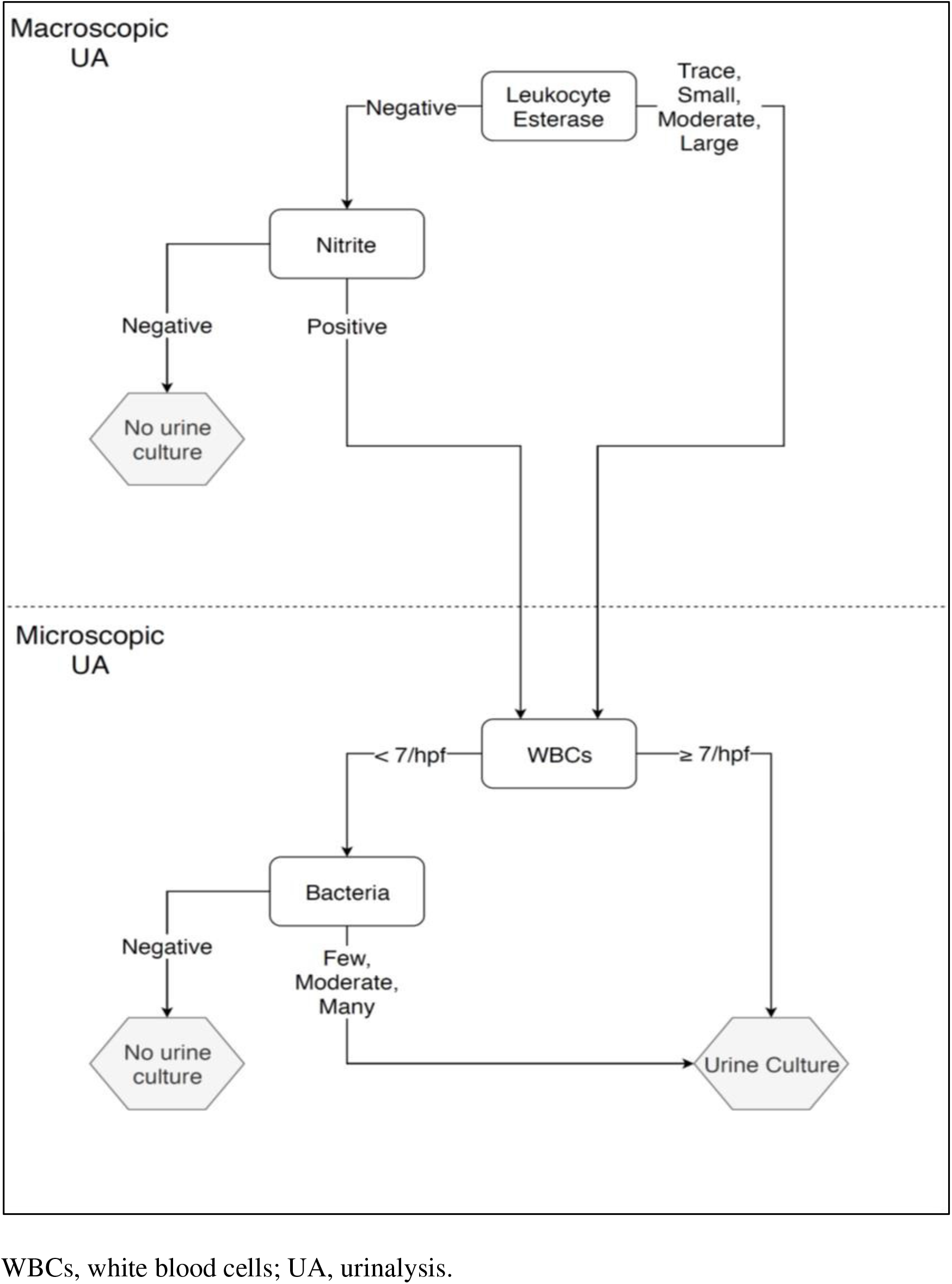
PittUDT algorithm decision tree for prediction of urine culture results using urinalysis parameters.

The test data comprised 9,773 cases, of which 2,571 (26.3%) had a positive UC result. Using the held-out test data set, the PittUDT algorithm met the pre-specified targets of a 30-60% total negative proportion (true negative plus false negative predictions) with a negative predictive value above 90% and a FNP below 5% for all hospitals (**Table 3**). Algorithm performance in the test data set after stratification by age (<60 years versus ≥60 years), gender, and specimen type is shown in **Table 2** and performance of the algorithm in the different settings (inpatient, outpatient, emergency department, and intensive care unit) are shown in **Table 3**.

**Table 3.** Summary of test data set PittUDT algorithm decision tree performance by hospital and location of specimen collection

| Performance Metric (n = 9,773) | Overall | Hospital 1 | Hospital 2 | Hospital 3 | Hospital 4 | Hospital 5 | Inpatient | Outpatient | ED | ICU |
| --- | --- | --- | --- | --- | --- | --- | --- | --- | --- | --- |
| Requires micro UA, n (%) | 5,673 (58) | 629 (78.4) | 1,166 (53.1) | 1,783 (62.6) | 1,190 (48.6) | 905 (61.1) | 1,246 (49.7) | 1,465 (48.4) | 2,698 (71.8) | 264 (55.2) |
| Requires UC, n (%) | 4,579 (46.9) | 525 (65.5) | 952 (43.4) | 1,380 (48.5) | 995 (40.6) | 727 (49.1) | 976 (38.9) | 1,132 (37.4) | 2,241 (59.6) | 230 (48.1) |
| TP, N (%) | 2,119 (21.7) | 278 (34.7) | 357 (16.3) | 757 (26.6) | 396 (16.2) | 331 (22.3) | 349 (13.9) | 513 (16.9) | 1,181 (31.4) | 76 (15.9) |
| TN, N (%) | 4,742 (48.5) | 251 (31.3) | 1,139 (51.9) | 1,328 (46.6) | 1,337 (54.6) | 687 (46.4) | 1,396 (55.7) | 1,765 (58.3) | 1,349 (35.9) | 232 (48.5) |
| FP, N (%) | 2,460 (25.2) | 247 (30.8) | 595 (27.1) | 623 (21.9) | 599 (24.5) | 396 (26.7) | 627 (25) | 619 (20.4) | 1,060 (28.2) | 154 (32.2) |
| FN, N (%) | 452 (4.6) | 26 (3.2) | 104 (4.7) | 139 (4.9) | 116 (4.7) | 67 (4.5) | 134 (5.3) | 132 (4.4) | 170 (4.5) | 16 (3.3) |
| TNP (%) | 53.1 | 34.5 | 56.6 | 51.5 | 59.4 | 50.9 | 61.1 | 62.6 | 40.4 | 51.9 |
| Sensitivity, % (95% CI) | 82.4 (80.9 - 83.9) | 91.4 (87.7 - 94.3) | 77.4 (73.3 - 81.2) | 84.5 (81.9 - 86.8) | 77.3 (73.5 - 80.9) | 83.2 (79.1 - 86.7) | 72.3 (68 - 76.2) | 79.5 (76.2 - 82.6) | 87.4 (85.5 - 89.1) | 82.6 (73.3 - 89.7) |
| Specificity, % (95% CI) | 65.8 (64.7 - 66.9) | 50.4 (45.9 - 54.9) | 65.7 (63.4 - 67.9) | 68.1 (65.9 - 70.1) | 69.1 (66.9 - 71.1) | 63.4 (60.5 - 66.3) | 69 (66.9 - 71) | 74 (72.2 - 75.8) | 56 (54 - 58) | 60.1 (55 - 65) |
| PPV, % (95% CI) | 46.3 (44.8 - 47.7) | 53 (48.6 - 57.3) | 37.5 (34.4 - 40.7) | 54.9 (52.2 - 57.5) | 39.8 (36.7 - 42.9) | 45.5 (41.9 - 49.2) | 35.8 (32.7 - 38.9) | 45.3 (42.4 - 48.3) | 52.7 (50.6 - 54.8) | 33 (27 - 39.5) |
| NPV, % (95% CI) | 91.3 (90.5 - 92.1) | 90.6 (86.5 - 93.8) | 91.6 (90 - 93.1) | 90.5 (88.9 - 92) | 92 (90.5 - 93.4) | 91.1 (88.9 - 93) | 91.2 (89.7 - 92.6) | 93 (91.8 - 94.1) | 88.8 (87.1 - 90.4) | 93.5 (89.7 - 96.3) |
| NA, % (95% CI) | 70.2 (69.3 - 71.1) | 66 (62.6 - 69.2) | 68.2 (66.2 - 70.1) | 73.2 (71.6 - 74.9) | 70.8 (68.9 - 72.6) | 68.7 (66.3 - 71.1) | 69.6 (67.8 - 71.4) | 75.2 (73.6 - 76.7) | 67.3 (65.8 - 68.8) | 64.4 (60 - 68.7) |
| BA, % (95% CI) | 74.1 (70.3 - 72) | 70.9 (57.4 - 61.6) | 71.6 (67.2 - 71.5) | 76.3 (70.9 - 73.9) | 73.2 (70.5 - 74.4) | 73.3 (67.3 - 71.7) | 70.6 (67.6 - 71.7) | 76.8 (73 - 76.5) | 71.7 (65.5 - 67.9) | 71.4 (63.9 - 71.9) |

BA, balanced accuracy; CI, confidence interval; ED, emergency department; FN, false negatives; FP, false positives; ICU, intensive care unit; TN, true negatives; TNP, total predicted negative proportion; TP, true positives; micro UA, microscopic urinalysis; UC, urine culture.

### Quality control (QC) program simulation

Data from all cases with complete data tested between October 1, 2017 and December 31, 2017 were used to simulate implementation of the validated algorithm at all five sites (n = 9,077 cases, of which 2,408 or 26.5% were UC positive). The results on one simulated QC run demonstrating four or fewer QC failures/month across all sites are provided in the supplemental materials (**Figure S1**); the actual FNP for each period is also presented. When the QC program was simulated 1,000 times, the QC testing failure rate was highest for sites with the highest actual FNP for that period (1.5% of simulations for site 1, 4.1% of simulations for site 2, 9.4% of simulations for site 3, 4.5% of simulations for site 4 and 0.4% of simulations for site 5).

### Standardization of laboratory procedures and electronic reporting

The SOPs for urinalysis and urine culture, reporting in the LIS, and middleware rules demonstrated differences across individual sites, which only became apparent when the processes for each site were reviewed together. To address inconsistencies between laboratories and to simplify ongoing evaluation, monitoring, and maintenance of the reflex testing procedure, a single SOP was created with a prescribed set of LIS and middleware rules to be enacted prior to implementation of the reflex testing algorithm for clinical use. The SOP was also developed to simplify the process of adopting the algorithm at additional sites in the future.

### Algorithm performance in real-world use

The performance of the PittUDT Urine Infection Testing algorithm across the 5 hospital sites was examined. A total of 594 randomly selected specimens (3 per day per site) which were classified by the algorithm as low risk with the UC not indicated were cultured. Of these, 589 of the specimens had a negative culture result. Across the 5 hospital sites, the concordance of the UC results predicted as low risk (UC negative) ranged from 98.4% to 100% (**Table 4**).

**TABLE 4.** Quality control summary by hospital

| **Facility** | **Date of Reflex Implementation** | **Number of Quality Control Samples** | **Negative Culture on Quality Control** | **(%)** |
| --- | --- | --- | --- | --- |
| Hospital 3 | 01/11/2021 | 400 | 397 | (99.3%) |
| Hospital 1 | 05/10/2021 | 113 | 112 | (99.1%) |
| Hospitals 2 and 4 | 09/26/2022 | 60 | 59 | (98.4%) |
| Hospital 5 | 09/26/2022 | 21 | 21 | (100%) |
| Total | - | 594 | 589 | (99.2%) |

## DISCUSSION

The implementation of a multi-site conditional UA reflex to UC protocol has the capacity to significantly decrease the number of unnecessary UC which are performed. UA parameters and thresholds used for reflex UC protocols vary widely and have historically been determined based on expert opinion or univariate ROC analysis.^15^ Our study demonstrates the feasibility and performance of using a supervised machine learning approach to develop a human readable machine learning model to optimize UA parameters in a reflex protocol that can be deployed across multiple hospital sites and settings within a large academic healthcare system. Our study also underscores the necessity for a stringent SOP review and robust data cleansing process performed by individuals with a granular understanding of the laboratory processes, middleware rules, and LIS rules, even when the laboratory results are generated at individual laboratories using the same instruments within one health system. Our study shows that a rule-based algorithm trained on UA/UC data has adequate predictive ability for triaging urine specimens by identifying those which are low risk and are unlikely to grow pathogenic organisms, with a false-negative proportion under 5%.

The PittUDT algorithm had similar performance in men and women, across an age range of adults, in different healthcare settings (inpatient, outpatient, emergency department, and intensive care unit), and for most specimen types (clean catch, straight catheter, and foley), except for suprapubic catheter and loop/ nephrostomy specimens. The decreased performance of the reflex algorithm with these cases may be due to the small sample size in the training and test data sets for these specimen types. However, it is also possible that UA is not an appropriate method of screening urine specimens obtained via suprapubic catheters and loop ileostomies or nephrostomies as patients with indwelling or suprapubic catheters and nephrostomy tubes invariably become carriers of asymptomatic bacteriuria, in which case antibiotic treatment does not have a benefit.^16,17^ In contrast to other studies on the use of artificial intelligence/machine learning (AI/ML) strategies for reducing unnecessary UC which specifically included special subpopulations of pregnant patients and children, our algorithm is not meant to be applied to these special groups.^18^ There was no uniform reliable mechanism to identify pregnant patients in the LIS, and although maternity-specific locations were excluded, it is unknown which patients in the dataset may have been pregnant. Furthermore, the guidance on these special groups is different from the general adult population. For example, the US Preventive Services Task Force recommends screening pregnant patients for asymptomatic bacteriuria, but not for nonpregnant adults.^19^

Other studies have developed ML algorithms for specific subpopulations or healthcare location settings. Taylor et al. applied supervised ML algorithms to predict UTIs in symptomatic patients in the ED, and Burton et al. applied supervised ML algorithms to predict UTIs in both inpatient and outpatient settings with laboratory testing performed in a single clinical microbiology laboratory covering three hospitals and community services.^18,20^ In contrast, we developed and deployed an ML algorithm which has broad application (inpatient, outpatient, emergency department, and intensive care unit) and demonstrated that strong clinical performance can be achieved using results from multiple laboratories by employing comprehensive preprocessing steps and a domain expertise level of understanding about what types of laboratory data are truly equivalent and can be combined.

The PittUDT algorithm was created using recursive partitioning to create a human readable decision tree by an expert group of laboratorians in conjunction with the Health System Infection Control Committee. This domain expertise was used to determine appropriate weights for false negatives and positives, and which metrics should be prioritized (e.g., negative predictive value). Our experience developing this algorithm underscores the importance of the laboratory director’s role in ensuring data integrity and equivalency in all of the nuanced decisions that form a foundational part of any ML algorithm implementation.

Other groups have evaluated other more complex ML methods for creating an algorithm for identifying low risk UA specimens, which are opaque to the end-user. Taylor et al. developed six ML models and concluded that of the ML models they evaluated, the XGBoost algorithm had the best performance.^20^ In contrast, Burton et al. found that when applied to their cohort, the sensitivity of the XGBoost algorithm was poor (61.7%) and hypothesized that the difference in performance (sensitivity) could be a consequence of the application of different class weights, as Burton et al. applied class weights that favored high sensitivity (desirable in a screening test) and Taylor et al. did not indicate if weighted classes were used.^18^

The development of ML algorithms for use in the clinical setting and evaluation of their performance cannot be relegated to computer scientists working in isolation; it requires clinical expertise and domain knowledge to understand whether appropriate decisions are made in both the selection and implementation of a particular model (e.g., appropriate population selection, equivalence or nonequivalence of laboratory data, understanding the cost of a false negative or positive test, and appropriate tuning such as class weighting). Furthermore, unlike other studies where all clinical testing was performed at a single laboratory, the macroscopic and microscopic UA and microbiology testing in our study was performed at multiple laboratory sites, both in the community hospital and large academic hospital settings. Our study also reveals a cautionary tale for those seeking to exclude laboratorians from these processes and simply combine laboratory data generated across multiple sites within a healthcare system or between healthcare systems based on the assumption that data generated for the same test, even when performed on the same instrument, should be equivalent. The preprocessing steps, evaluation of the SOPs, middleware and LIS rules, were all integral to developing a sound dataset prior to algorithm development.

While the PittUDT algorithm can be finetuned for differences in the hospitals, locations within a hospital, and sex, the goal of this study was to create a universal algorithm for all sites utilizing the Sysmex UF-1000i (Sysmex, Lincolnshire, IL) and Clinitek Atlas or Clinitek Novus (Clinitek, Ramsey, MN) urinalysis instruments. As documented in the literature, urine WBCs and bacteria were the best predictors of positive UC.^21^ These two parameters are derived from the automated microscopic analysis, which has high reproducibility and accuracy especially when performed by flow cytometry.^22^ Urine leukocyte esterase, which is a semi-quantitative measure of the number of WBCs, was also a good predictor of UC positivity. A universal algorithm based on data driven parameters with acceptable performance across a diversity of patients is a formidable but attainable achievement which facilitates more rapid implementation and change control in a multisite healthcare system.

