## Supplemental Tables 1 and 2, Supplemental Figure 1 for "Development and Evaluation of a Machine Learning Recursive Partitioning Decision Tree Algorithm to Optimize Urinalysis Parameters to Predict Urine Culture Positivity"

**SUPPLEMENT MATERIAL**

**Tables**

**Table S1.** Summary data for results of macroscopic urinalysis for all cases by hospital

| **Site** |  | **Hospital 1** | **Hospital 2** | **Hospital 3** | **Hospital 4** | **Hospital 5** | **Overall** |
| --- | --- | --- | --- | --- | --- | --- | --- |
| Leukocyte Esterase, n (%)^a^ | Negative | 795 (23.7) | 4,034 (48.4) | 4,317 (38.9) | 5,003 (53) | 2,432 (39.7) | 16,581 (43.2) |
|  | Trace | 596 (17.8) | 781 (9.4) | 1,464 (13.2) | 786 (8.3) | 702 (11.5) | 4,329 (11.3) |
|  | Small | 514 (15.3) | 1,416 (17) | 2,213 (20) | 1,350 (14.3) | 1,177 (19.2) | 6,670 (17.4) |
|  | Moderate | 570 (17.0) | 1,136 (13.6) | 1,594 (14.4) | 1,188 (12.6) | 818 (13.4) | 5,306 (13.8) |
|  | Large | 881 (26.3) | 975 (11.7) | 1,505 (13.6) | 1,121 (11.9) | 993 (16.2) | 5,475 (14.3) |
| Nitrite, n (%)^a^ | Negative | 2,681 (79.9) | 7,580 (90.9) | 9,617 (86.7) | 8,393 (88.8) | 5,152 (84.2) | 33,423 (87.1) |
|  | Positive | 675 (20.1) | 762 (9.1) | 1,476 (13.3) | 1,055 (11.2) | 970 (15.8) | 4,938 (12.9) |
| Protein, n (%)^a^ | Negative | 1,570 (46.8) | 5,637 (67.6) | 5,807 (52.4) | 4,986 (52.8) | 2,440 (39.9) | 20,440 (53.3) |
|  | Trace | 418 (12.5) | 1,021 (12.2) | 1,695 (15.3) | 1,277 (13.5) | 1,073 (17.5) | 5,484 (14.3) |
|  | Small | 608 (18.1) | 906 (10.9) | 1,879 (16.9) | 1,595 (16.9) | 1,346 (22) | 6,334 (16.5) |
|  | Moderate | 497 (14.8) | 586 (7) | 1,362 (12.3) | 1,199 (12.7) | 999 (16.3) | 4,643 (12.1) |
|  | Large | 263 (7.8) | 192 (2.3) | 350 (3.2) | 391 (4.1) | 264 (4.3) | 1,460 (3.8) |
| Blood, n (%)^a^ | Negative | 1,408 (42.0) | 4,750 (56.9) | 5,999 (54.1) | 4,591 (48.6) | 2,738 (44.7) | 19,486 (50.8) |
|  | Trace | 483 (14.4) | 1,028 (12.3) | 1,235 (11.1) | 1,224 (13) | 700 (11.4) | 4,670 (12.2) |
|  | Small | 381 (11.4) | 743 (8.9) | 1,089 (9.8) | 1,059 (11.2) | 774 (12.6) | 4,046 (10.6) |
|  | Moderate | 462 (13.8) | 723 (8.7) | 1,055 (9.5) | 1,017 (10.8) | 715 (11.7) | 3,972 (10.4) |
|  | Large | 622 (18.5) | 1,098 (13.2) | 1,715 (15.5) | 1,557 (16.5) | 1,195 (19.5) | 6,187 (16.1) |
| pH,  median (5^th^–95^th^ percentiles)  (range)^b^ |  | 6.0  (5.0 - 8.0;  5.0 - 9.0) | 6.0  (5.0 - 7.5;  5.0 - 9.0) | 6.0  (5.5 - 7.5;  5.0 - 9.0) | 6.0  (5.0 - 7.5;  5.0 - 9.0) | 6.0  (5.0 - 7.5;  5.0 - 8.5) | 6.0  (5.0 - 7.5;  5.0 - 9.0) |
| Specific gravity,  median (5^th^– 95^th^ percentiles)  (range)^b^ |  | 1.018  (1.005 - 1.030; 1.001 - 1.071) | 1.017  (1.005 - 1.033; 1.000 - 1.090) | 1.02  (1.008 - 1.031; 1.000 - 1.125) | 1.018  (1.007 - 1.035; 1.000 - 1.099) | 1.018  (1.007 - 1.035; 1.001 - 1.098) | 1.018  (1.006 - 1.033; 1.000 - 1.125) |
| Character, n (%)^a^ | Clear | 1,762 (52.5) | 4,701 (56.4) | 5,404 (48.7) | 5,660 (59.9) | 2,582 (42.2) | 20,109 (52.4) |
|  | Hazy | 55 (1.6) | 38 (0.5) | 1,018 (9.2) | 54 (0.6) | 13 (0.2) | 1,178 (3.1) |
|  | Cloudy | 1,536 (45.8) | 2,865 (34.3) | 3,615 (32.6) | 2,699 (28.6) | 2,637 (43.1) | 13,352 (34.8) |
|  | Turbid | 3 (0.1) | 738 (8.9) | 1,056 (9.5) | 1,035 (11) | 890 (14.5) | 3,722 (9.7) |

^a^Categorical data are summarized using frequency (%).

^b^Interval or ratio data are presented as the median (5^th^ – 95^th^ percentile) and (range).

**Table S2.** Summary data for results of microscopic urinalysis and urine culture positivity rate for all cases by hospital

| **Parameter** | **Hospital 1** | **Hospital 2** | **Hospital 3** | **Hospital 4** | **Hospital 5** | **Overall** |
| --- | --- | --- | --- | --- | --- | --- |
| Bacteria (cells/µL)^a^ | 0-500: 1,905 (56.8) | 0-600: 5,173 (62) | 0-500: 5,756 (51.9) | 0-600: 6,733 (71.3) | 0-500: 4,227 (69) |  |
|  | 500-1500: 365 (10.9) | 600-1200: 1,348 (16.2) | 500-1000: 1,003 (9) | 600-1200: 1,081 (11.4) | 500-1500: 540 (8.8) |  |
|  | 1500-3500: 273 (8.1) | 1200-2400: 710 (8.5) | 1000-3500: 2,193 (19.8) | 1200-2400: 453 (4.8) | 1500-3500: 294 (4.8) |  |
|  | 3500-5000: 115 (3.4) | >2400: 1,111 (13.3) | 3500-5000: 737 (6.6) | >2400: 1,181 (12.5) | 3500-5000: 113 (1.8) |  |
|  | >5000: 698  (20.8) |  | >5000: 1,404 (12.7) |  | >5000: 948 (15.5) |  |
| WBC (cells/hpf)  median (5^th^– 95^th^ percentiles)  (range)^b^ | 7  (3 - >49;  0 - >49) | 4  (0 - 229;  0 - 3,036) | 5  (0 - 324;  0 - 900) | 4  (0 - 252;  0 - 3,257) | 7  (1 - 616;  0 - 1,797) | 5  (0 - 286;  0 - 3,257) |
| RBC (cells/hpf)  median (5^th^– 95^th^ percentiles)  (range)^b^ | 1  (1 - >30;  0 - >30) | 3  (1 - 100;  0 - 6,236) | 3  (0 - 100;  0 - 9,378) | 4  (1 - 139;  0 - 9,772) | 5  (1 - 474;  0 - 8,438) | 3  (0 - 107;  0 - 9,772) |
| Positive UC, n (%)^a^ | 1,270 (37.8) | 1,744 (20.9) | 3,548 (32.0) | 2,037 (21.6) | 1,603 (26.2) | 10,202 (26.6) |

^a^Categorical data are summarized using frequency (%).

^b^Interval or ratio data are presented as the median (5^th^ – 95^th^ percentile) and (range).

Hpf, high power field; UC, urine culture.

**Figures**

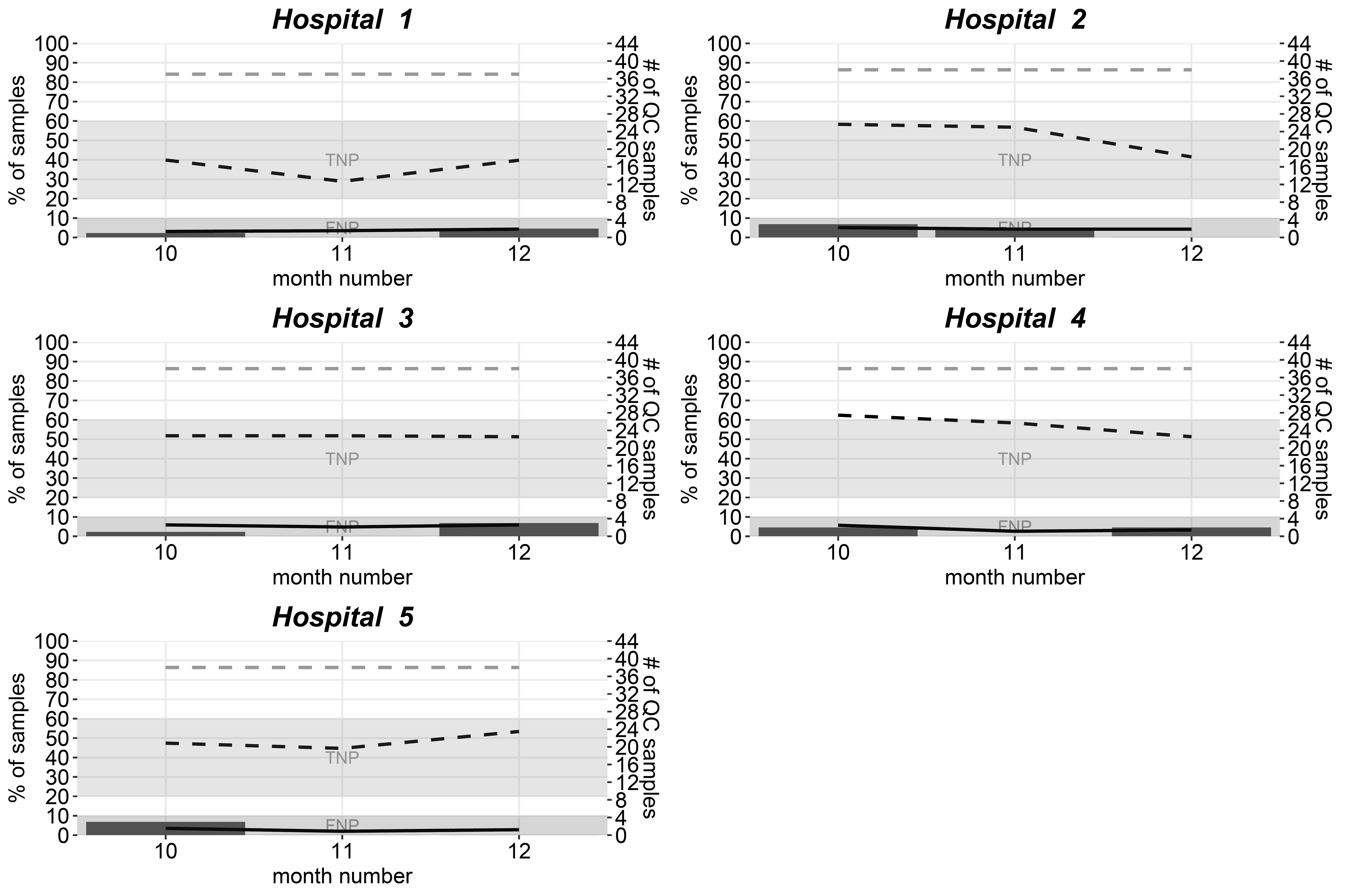

**Figure S1.** Simulated performance of quality control program based on two-stage hypergeometric sampling plan for all cases. Data between October 1, 2017 and December 31, 2017 were used to simulate implementation of the validated algorithm at all five sites (n = 9,077 cases, of which 2,408 or 26.5% were urine culture positive). The number of quality control samples tested each month is shown in the light grey dashed line, while the actual proportion of total negative predictions and false negative predictions per month are shown in the black dashed and solid lines, respectively. Control limits for the total negative proportion and false negative proportion are shaded. The number of quality control failures in the tested samples are shown as bars.
